# Increased thermostability of an engineered flavin-containing monooxygenase to remediate trimethylamine in fish protein hydrolysates

**DOI:** 10.1101/2023.03.10.532160

**Authors:** Marianne Goris, Isabel Cea-Rama, Pål Puntervoll, Rasmus Ree, David Almendral, Julia Sanz-Aparicio, Manuel Ferrer, Gro Elin Kjæreng Bjerga

## Abstract

Protein hydrolysates made from marine by-products are very nutritious, but frequently contain trimethylamine (TMA) which has an unattractive fish-like smell. Bacterial trimethylamine monooxygenases can oxidize TMA into the odorless trimethylamine *N*-oxide (TMAO) and have been shown to reduce TMA-levels in a salmon protein hydrolysate. To make the *Methylophaga aminisulfidivorans* trimethylamine monooxygenase, mFMO, more suitable for industrial application, we engineered it using the Protein Repair One-Stop Shop (PROSS) algorithm. All seven mutant variants, containing 8-28 mutations, displayed increases in melting temperature between 4.7 °C and 9.0 °C. The crystal structure of the most thermostable variant, mFMO_20, revealed the presence of four new stabilizing interhelical salt bridges, each involving a mutated residue. Finally, mFMO_20 significantly outperformed native mFMO in its ability to reduce TMA levels in a salmon protein hydrolysate at industrially relevant temperatures.

**Importance:** Marine by-products are a high-quality source for peptide ingredients, but the unpleasant fishy odour caused by TMA limits their access to the food market. This problem can be mitigated by enzymatic conversion of TMA into the odourless TMAO. Enzymes isolated from nature must be adapted to industrial requirements, however, such as the ability to tolerate high temperatures. This study has demonstrated that mFMO can be engineered to become more thermostable. Moreover, unlike the native enzyme, the best thermostable variant efficiently oxidized TMA in a salmon protein hydrolysate at industrial temperatures. Our results present an important next step towards application of this novel and highly promising enzyme technology in marine biorefineries.

## Introduction

Flavin containing monooxygenases (FMOs, EC 1.14.13.8) are enzymes that insert one molecule of oxygen into organic substrates using the cofactors FAD and NAD(P)H (Ceccoli et al., 2014; Torres Pazmiño et al., 2010; van Berkel et al., 2006). A subgroup of bacterial FMOs oxidize trimethylamine (TMA) to trimethylamine *N*-oxide (TMAO) and are often referred to as trimethylamine monooxygenases (Tmms) (Chen et al., 2011; Choi et al., 2003; Goris et al., 2020). These and other FMOs have also gained interest for their ability to convert indole into the dye indigo and the drug agent indirubin (Choi et al., 2003; Fabara and Fraaije, 2020; Han et al., 2012; Rioz-Martínez et al., 2011).

TMA is a well-known contributor to the odor of spoiled fish (Hebard et al., 1982), and may accumulate to give rise to a strong bodily odor in humans with trimethylaminuria (fish odor syndrome) caused by impairments in the *FMO3* gene (Schmidt and Leroux, 2020).

Fish protein hydrolysates made from by-products from fisheries and aquaculture are of high nutritional value, and have a great potential for the human consumption market (Shavandi et al., 2019; Villamil et al., 2017). However, fish protein hydrolysates frequently suffer from an off-putting malodor which is mainly caused by TMA. Currently, the TMA malodor may be handled by odor masking, vaporization, encapsulation or filtration, albeit with varying degrees of success and possibly also compromising other qualities in the products. Application of Tmm enzymes is thus an alternative and novel strategy to convert TMA to the odorless TMAO in fish protein hydrolysates. This has the potential to significantly improve the organoleptic quality of fish protein hydrolysates and thereby promoting their application as food ingredients, while simultaneously maintaining their nutritional profile.

In a previous study, we screened 45 bacterial Tmms for their ability to oxidize TMA to TMAO (Goris et al., 2020), and identified the *Methylophaga aminisulfidivorans* Tmm (mFMO) (Choi et al., 2003) as a suitable candidate for application on a TMA-containing salmon protein hydrolysate. In industrial fish protein hydrolysis, enzymes are required to perform at pH around 6 and temperatures ranging from 45 °C to 60 °C (Aspevik et al., 2016; Kristinsson and Rasco, 2000). This implies that mFMO, with an optimal temperature of 44.0 °C and melting temperature of 46.7 °C, would benefit from enzyme engineering to increase its stability (Goris et al., 2020). In that respect, a previous effort to engineer mFMO is encouraging. Lončar and colleagues used the computational protocol FRESCO (Lončar et al., 2019; Wijma et al., 2018) to predict two mutations in a mFMO, M15L and S23A, that when combined increased the apparent melting temperature by 3.0 °C (Lončar et al., 2019).

The mFMO enzyme forms a dimer and each monomer consists of two domains: the larger FAD-binding domain and the smaller NADPH-binding domain (Alfieri et al., 2008; Cho et al., 2011). Upon binding, NADPH reduces the tightly bound FAD, thus generating the reactive flavin intermediate C4a-hydroperoxy-FAD and NADP^+^. The latter stabilizes the activated flavin intermediate, and together with residue tyrosine 207, it shields the active site and the intermediate from the solvent (Alfieri et al., 2008). When entering the active site, the substrate displaces NADP^+^ and is subsequently oxidized by the activated flavin intermediate.

Protein Repair One-Stop Shop (PROSS) is a web server that takes a protein structure as input and outputs several mutated sequences that are expected to have increased stability (Goldenzweig et al., 2016). PROSS combines multiple independently stabilizing mutations by integrating Rosetta modelling and phylogenetic sequence information (Goldenzweig et al., 2016). In a recent community-wide experimental evaluation of PROSS, designs for nine of ten tested protein targets displayed increased temperature stability, ranging from 8.3 °C to 27.0 °C (Peleg et al., 2021).

In the present study, we employed the PROSS algorithm on mFMO to improve its thermal stability. Seven combinatorial mutant variants of mFMO, containing 8-28 mutations, were analyzed for their temperature stability and compared to wild type mFMO. We demonstrate that all mFMO variants were more thermostable than the wild type. The most thermostable variant was analysed by steady state kinetics and compared to the wild type without identifying substantial modification of the kinetic parameters.

Moreover, this stabilized mFMO variant also converted TMA to TMAO more efficiently than native mFMO in a salmon protein hydrolysate at two industrially relevant temperatures, 50.0 °C and 65.0 °C. Finally, the crystal structure of the most thermostable variant was solved to elucidate the structural basis for the increased thermal stability, revealing loss of flexibility through a novel network of polar interactions as the main contributing factor.

## Results

### Design and expression of mutant variants of mFMO with predicted increased stability

To make a more stable and temperature resistant mFMO, ideally withstanding at least 50 °C in industrial applications, we employed computational enzyme engineering. The most recent crystal structure of mFMO in complex with the cofactors FAD and NADP^+^ (PDB ID:2XVH) (Cho et al., 2011) was used as input to the PROSS web server (Goldenzweig et al., 2016) along with instructions to exclude residues in contact with the cofactors, as well as dimer interface residues, as mutational targets. PROSS proposed 7 mFMO variants with the number of mutations ranging from 8 to 28 (Figure 1, Figure S1). The variants were named mFMO_*n*, where *n* indicates the number of mutations. The mutations were located at or near the surface, and the number of residues predicted to form new stabilizing salt bridges increased from 2 in mFMO_8 to 8 in mFMO_28 (Figure 1, Table 1). In the models of mFMO_8 through mFMO_20, all new salt bridges were predicted to form between one mutated and one native residue (Table 1). The last two variants displayed more complex salt bridge patterns: In mFMO_24, the newly introduced N394K mutation is predicted to form salt bridges with both T370D and D374, and in mFMO_28, the newly introduced P391D mutant is predicted to form a third salt bridge with N394K (Table 1). Interestingly, the majority of the new salt bridges are predicted to form interhelical connections (Table 2).

**Figure 1.**
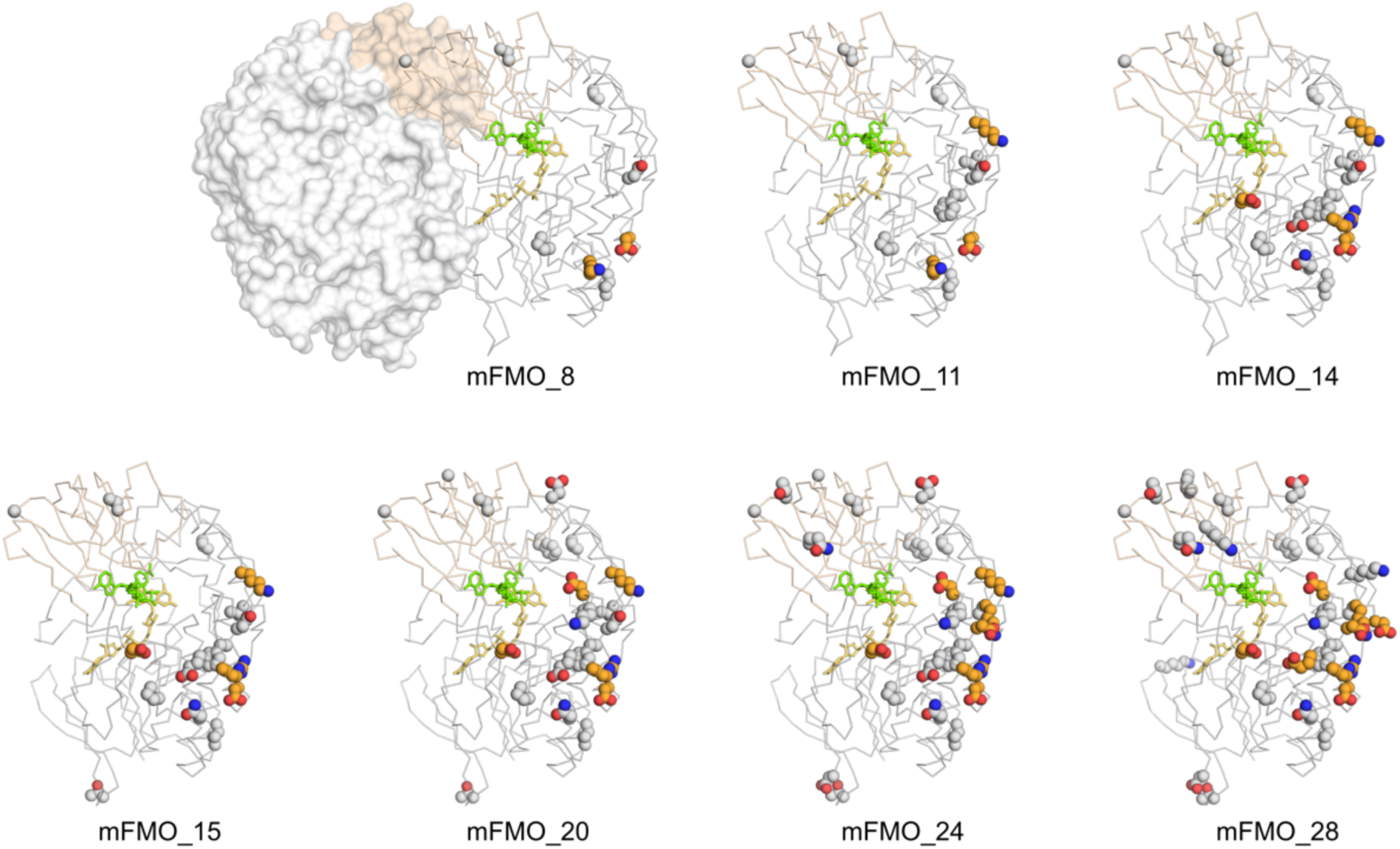
Structural models of mFMO and mutant variants. The structural models of the 7 variants of mFMO with 8 to 28 mutations (top left to bottom right), as generated by PROSS (using PDB:2XVH), are shown in ribbon view (chain A only). For context and to indicate the dimer interface, chain B of native mFMO is shown in surface view (top left, PDB:2XVH). The large domain is shown in grey, the small domain in wheat, and the cofactors FAD and NADP^+^ are shown in yellow and green sticks, respectively. Mutant residues predicted to form salt bridges are shown as orange spheres, and other mutations are shown as grey spheres. Nitrogen and oxygen atoms of the mutant residues are colored blue and red, respectively.

**Table 1.** Predicted new salt bridges formed between mutated residues and native or mutated residues. Empty cells indicate the absence of the mutated residue listed to the left. Residues (native or mutant) forming a predicted salt bridge with the mutant residue (left column) are listed, and if a mutant residue does not form a salt bridge in an mFMO variant, it is marked by a minus (-). The total number of mutant residues involved in forming salt bridges are shown in the bottom row.

| Mutated residue | Mutant mFMO variant |  |  |  |  |  |  |
| --- | --- | --- | --- | --- | --- | --- | --- |
|  | mFMO_8 | mFMO_11 | mFMO_14 | mFMO_15 | mFMO_20 | mFMO_24 | mFMO_28 |
| N290D |  |  | R292 | R292 | R292 | R292 | R292 |
| M353K | E357 | E357 |  |  |  |  |  |
| K358R |  |  | D374 | D374 | D374 | D374 | D374 |
| L360E |  |  | - | - | - | - | R356/R292 |
| A365D |  |  |  |  | K401 | K401 | K401 |
| T370D | - | - | - | - | - | N394K | N394K |
| N378D | K300 | K300 | K300 | K300 | K300 | K300 | K300 |
| P391D |  |  |  |  |  |  | N394K |
| N394K |  |  |  |  |  | T370D/D374 | T370D/D374/P391D |
| L398K |  | E366 | E366 | E366 | E366 | E366 | - |
| **Total** | **2** | **3** | **4** | **4** | **5** | **7** | **8** |

**Table 2.** Predicted interhelical salt bridges involving or induced by mutated residues in PROSS mutant models. Note that the α6-β18/β19-α7 salt bridge connects helices α6 and α7 via the β18/β19 loop and that the α6-β17/β18 salt bridge connects helix α6 to the β17/β18 loop.

| mFMO variant | Interhelical salt bridges |  |  |  | Total |
| --- | --- | --- | --- | --- | --- |
| | $\alpha 6$-$\alpha 7$ | $\alpha 6$-$\beta 18/\beta 19$-$\alpha 7$ | $\alpha 7$-$\alpha 8$ | $\alpha 6$-$\beta 17/\beta 18$ | |
| mFMO_8 |  | D351-K300-N378D |  |  | 1 |
| mFMO_11 |  | D351-K300-N378D | E366-L398K |  | 2 |
| mFMO_14 | K358R-D374 | D351-K300-N378D | E366-L398K |  | 3 |
| mFMO_15 | K358R-D374 | D351-K300-N378D | E366-L398K |  | 3 |
| mFMO_20 | K358R-D374 | D351-K300-N378D | E366-L398K; A365D-K401 |  | 4 |
| mFMO_24 | K358R-D374 | D351-K300-N378D | E366-L398K; A365D-K401; T370D-N394K; D374-N394K |  | 6 |
| mFMO_28 | K358R-D374 | D351-K300-N378D | A365D-K401; T370D-N394K; D374-N394K | L360E-R292 | 6 |

All 7 mFMO mutant variants were expressed with a C-terminal hexa-histidine tag, at levels comparable to that of native mFMO, purified (Figure S2), and verified by mass spectrometry. When expressing mFMO in *Escherichia coli*, the culture medium turns blue due to the enzymatic conversion of endogenous indole to indigo (Choi et al., 2003; Fabara and Fraaije, 2020; Goris et al., 2020). The fact that the culture media of all mFMO variants turned blue following overnight expression suggested that the expressed mFMO variants were functional. Moreover, all purified mFMO variant enzymes were colored bright yellow, indicating the presence of bound FAD cofactor, which is required for function.

### mFMO mutant variants are functional and more thermostable than native mFMO

To investigate whether the mFMO variants had increased thermal stability compared to the native enzyme, we conducted protein melting studies using circular dichroism. The melting temperature (*T*_m_) of native mFMO was measured to be 46.2 °C (Table 3), which is in line with previous results (Goris et al., 2020). All mFMO variants demonstrated increased temperature stability compared to the wildtype enzyme, as reflected by their *T*_m_ values, which ranged from 50.9 °C for mFMO_28 to 55.2 °C for mFMO_20 (Table 3). The melting temperature increased with the number of mutations from mFMO_8 to mFMO_20 but declined slightly for mFMO_24 and mFMO_28.

**Table 3.** Biochemical parameters of mFMO and mutant variants. Melting curves were obtained by CD at pH 7.5 and the melting temperature (*T*_m_ ± 95% CI) was estimated by four-parameter logistic regression of the melting curve. Temperature where half enzyme activity is lost (*T_1/2_*) was measured at pH 7.5 and the reported values are means of three independent experiments (± SD). Optimal temperature (*T*_opt_) was determined at pH 8.0 for mFMO and mFMO_20. Optimal pH for TMA conversion was determined at 22 °C. Steady-state kinetic measurements were performed with varying concentrations of TMA (Sigma-Aldrich) as the substrate with fixed NADPH concentration (200 µM) at 23°C, pH 8.0. Reported values are the means of two biological replicates (± SD).

| Enzyme | $T_m$ (°C) | $T_{1/2}$ (°C) | $T_{opt}$ (°C) | Optimal pH | $K_M$ (μM) | $k_{cat}$ ($s^{-1}$) | $k_{cat}/K_M$ (μM$^{-1}$ $s^{-1}$) |
| --- | --- | --- | --- | --- | --- | --- | --- |
| mFMO | 46.2±0.2 | 40.6±0.4 | 40 | 8.5 | 1.07±0.13 | 1.28±0.03 | 1.20±0.12 |
| mFMO_8 | 51.2±0.1 | 45.1±0.9 | - | 7.5 | - | - | - |
| mFMO_11 | 51.7±0.1 | 47.1±1.2 | - | 7.5 | - | - | - |
| mFMO_14 | 53.9±0.2 | 49.0±1.5 | - | 8.0 | - | - | - |
| mFMO_15 | 54.5±0.1 | 49.6±0.4 | - | 7.5 | - | - | - |
| mFMO_20 | 55.2±0.1 | 50.0±0.4 | 40 | 8.0 | 0.83±0.02 | 0.93±0.05 | 1.11±0.03 |
| mFMO_24 | 52.0±0.1 | 49.2±0.2 | - | 7.5 | - | - | - |
| mFMO_28 | 50.9±0.1 | 48.9±0.5 | - | 8.0 | - | - | - |

To study the increased temperature stability of the mFMO variants further, we evaluated their remaining catalytic activity against TMA after one-hour incubations at temperatures from 30.0 °C to 54.0 °C. The temperature at which half the enzyme activity was lost (*T*_1/2_) ranged from 45.1 °C for mFMO_8 to 50.0 °C for mFMO_20, all outperforming native mFMO which had a *T*_1/2_ of 40.6 °C (Table 3). The *T*_1/2_ values increased with the number of mutations in the same manner as the *T*_m_, with a moderate decline recorded for mFMO_24 and mFMO_28.

As production of protein hydrolysates is often performed between pH 6.0 and 7.0 (Aspevik et al., 2016; Kristinsson and Rasco, 2000), we also assessed whether engineering altered the pH optimum, which was previously determined to be 8.5 for native mFMO (Goris et al., 2020). As seen in Table 3, all mFMO variants had pH optima between 7.5 and 8.0, which are slightly lower than that of native mFMO.

Although mFMO_20 did not have the lowest pH optimum among the mutant variants, it displayed the greatest improvement in temperature stability, as reflected by both *T*_m_ and *T*_1/2_. As mFMO_20 thus emerged as the most promising variant for industrial application, we determined its optimal temperature (*T*_opt_) for enzymatic activity and compared it to that of native mFMO. To determine *T*_opt_, we assessed the specific activity against TMA at temperatures between 22 °C and 50 °C, at pH 8.0 (Figure S3). The *T*_opt_ for both native mFMO and mFMO_20 was found to be 40 °C (Table 3). The *T*_opt_ for native mFMO was previously reported to be 45 °C (at pH 7.5) (Goris et al., 2020), but the observed differences in the activity measured at 40 and 45 °C in both studies were marginal.

Engineering enzymes to increase stability often comes with a trade-off of diminished catalytic activity (Klesmith et al., 2017). We therefore performed a steady-state kinetic analysis of native mFMO and mFMO_20, using TMA as substrate with fixed concentrations of NADPH (Table 3). Under the conditions tested, the 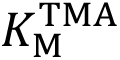 of native mFMO was 1.07 µM and the 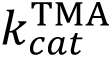 was 1.28 s^-1^. The 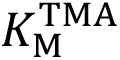 values of mFMO_20 was 0.83 µM and the 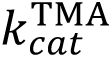 was 0.93 s^-1^, both slightly lower than that of native mFMO. Interestingly, the catalytic efficiency (*k*_cat_/*K*_M_) of mFMO_20 remained almost identical to that of native mFMO (Table 3).

### mFMO_20 reduces the TMA level by 95% in salmon protein hydrolysate at 65 °C

Since mFMO_20 demonstrated the most prominent increase in thermal stability, we compared its ability to convert TMA to TMAO in a salmon protein hydrolysate to that of native mFMO. The cofactor NADPH was supplemented, as the hydrolysate did not contain sufficient amounts to drive the enzymatic reaction (Goris et al., 2020). Heat-calibrated enzymes and 0.5 mM NADPH were added to salmon protein hydrolysates (pH 6.1) and incubated for 1 hour at 30 °C, 50 °C and 65 °C, followed by measurements of TMA and TMAO concentrations (Figure 2). When treated with the native enzyme, the TMA level in the hydrolysate was reduced by 52% at 30°C and 46% at 50°C, and only 29% reduction was observed at 65°C. The mutant variant mFMO_20 outperformed native mFMO at all temperature with a striking 95% reduction of TMA at both 50°C and 65°C.

**Figure 2.**
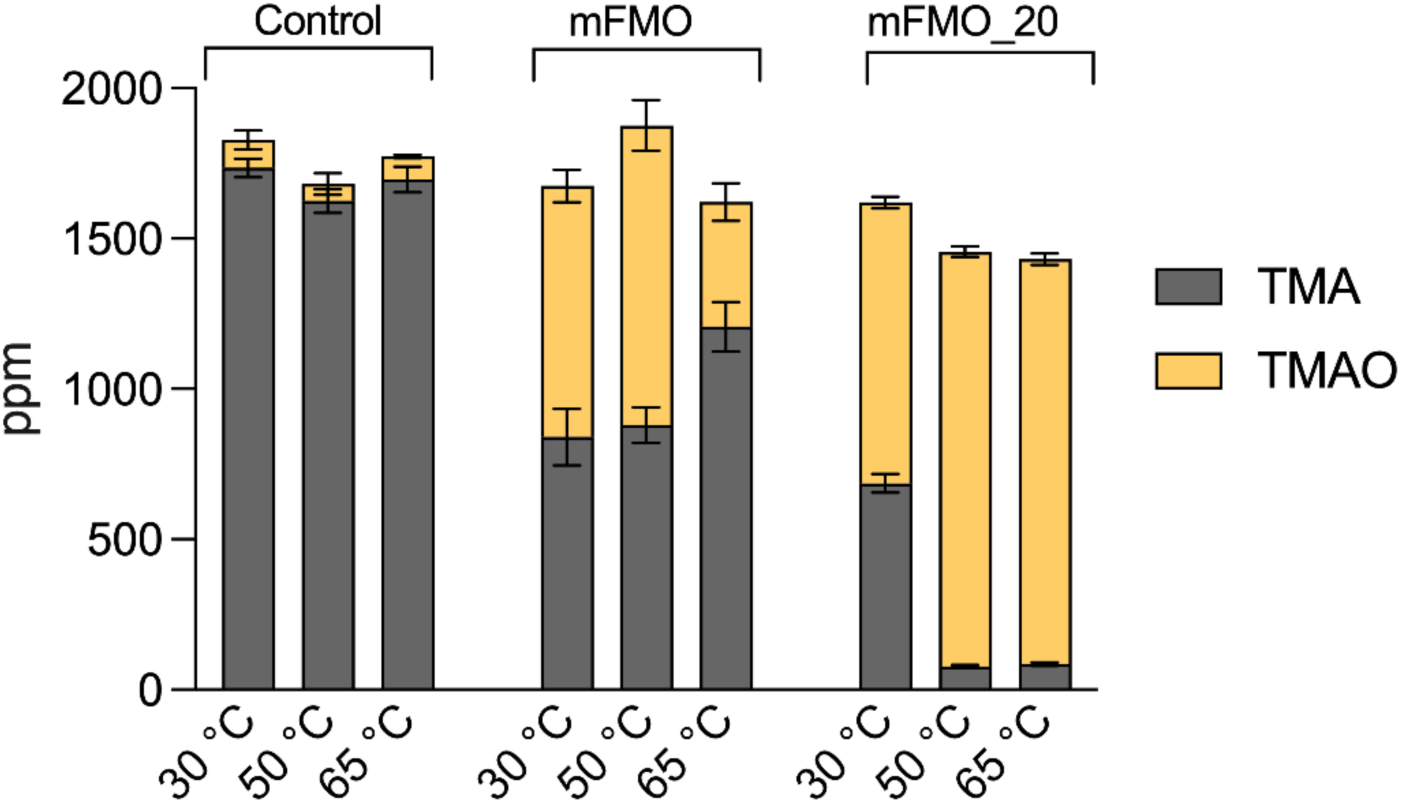
Enzymatic TMA conversion in salmon protein hydrolysate. Salmon protein hydrolysates (pH 6.1) were incubated for 1 hour at 30, 50 and 65 °C with no enzyme (control), mFMO, or mFMO_20, all supplemented with 0.5 mM NADPH. TMA and TMAO levels were determined using UHPLC with EVOQ Elite Triple Quadrupole Mass Spectrometer. The experiment was performed twice, each time with three technical replicates, and the plot shows the mean TMA (grey) and TMAO (yellow) levels (ppm) in stacks with standard deviations represented by error bars.

### Network of novel polar interactions stabilizes mFMO_20

To understand the structural basis for the increased thermostability of mFMO_20, we crystallized it with the cofactors FAD and NADPH. The crystals were indexed in the C222_1_ space group and contained the biological dimer within the asymmetric unit, with one FAD and one NADP^+^ molecule bound per catalytic site. The crystal structure of the mFMO_20/FAD/NADP^+^ complex was solved at 1.62 Å resolution, revealing a structure highly similar to that of native mFMO (PDB ID: 2XVH) (Cho et al., 2011), as reflected by a calculated RMSD of 0.27 Å (on 445 C*α*atoms). The small domain contains 4 mutations, and the large domain contains the other 16 (Figure 3). Compared to native mFMO, mFMO_20 has a net charge change of -4. Interestingly, half of the mFMO_20 mutations are located in a 46-amino-acid subsequence (M353Q-L398K) of a region in the large domain containing three helices: *α*6 (K345-T361), *α*7 (A365D-M382), and *α*8 (I390-N406) (Figure 3). Structural analysis revealed that five of these mutated residues form new salt-bridges involving six native residues (K358R-D374, L360E-R356, A365D-K401, N378D-K300-D351, and L398K-E366), of which four form interhelical interactions (Figure 4A, Table 4), thus confirming the PROSS model predictions (Table 2). The helices *α*6 and *α*7 are directly connected by K358R-D374, and indirectly connected, via the β18-β19 loop, by N378D-K300-D351. The helices *α*7 and *α*8 are connected by A365D-K401 and L398K-E366. Two new hydrogen bonds involving side chains were also introduced: one forming an intrahelical bond (*α*6; M353Q-R356) and the other an interhelical bond (*α*7-*α*8; T370D-N394) (Table 4). In addition to the salt bridges directly introduced by mutated residues, mFMO_20 has 4 new salt bridges involving native residues, compared to native mFMO (Figure 4A). All but one of the salt bridges identified in native mFMO were also present in mFMO_20 (Figure 4B).

**Figure 3.**
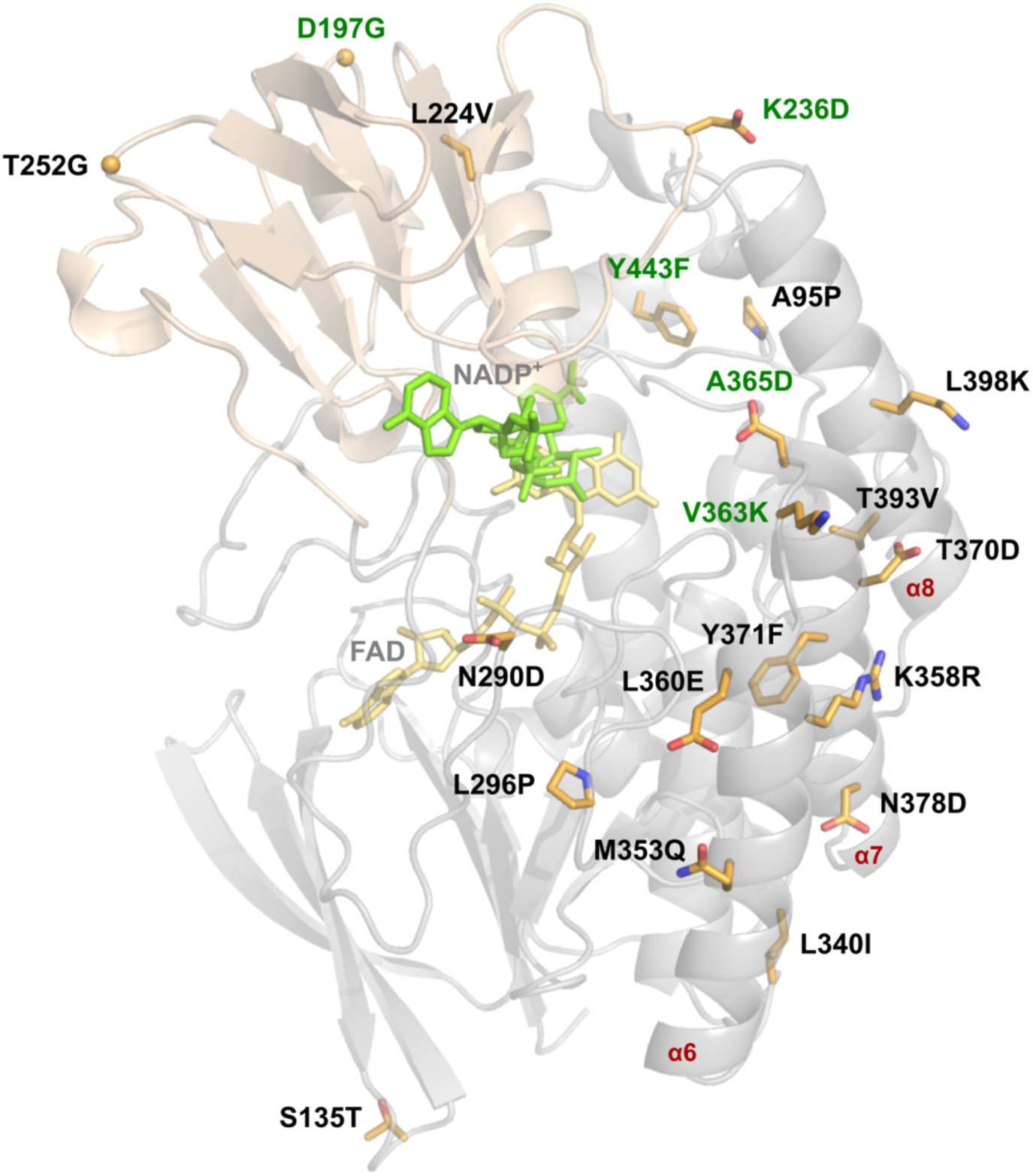
Crystal structure of mFMO_20. The structure of the mFMO_20 monomer (PDB ID: 8B2D) (chain A) is shown as cartoon, with the large domain colored in gray and the small domain colored in wheat. The cofactors FAD and NADP^+^ are shown as yellow and green sticks, respectively. The side chains of the 20 mutated residues are represented as orange sticks, and oxygen and nitrogen atoms colored red and blue, respectively. The alpha carbons of the introduced glycine residues are shown as spheres. The 5 residues that are new compared to mFMO_15 are labeled in green. The three α-helices where the 10 of 20 mutated residues are located are labeled in red.

**Figure 4.**
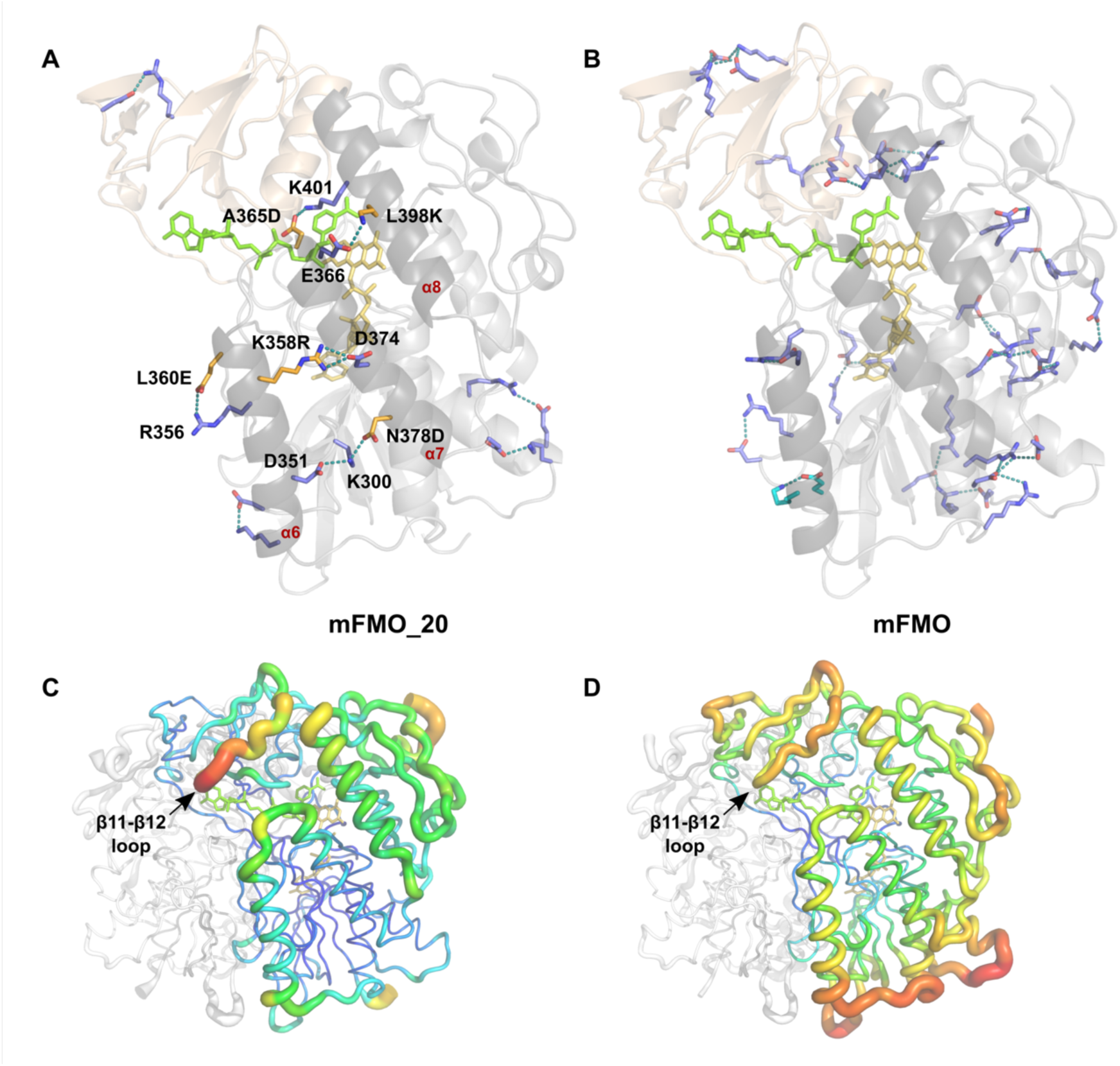
Stabilizing salt bridges in native mFMO and mutant mFMO_20. **A)** Chain A of the mFMO_20 structure is shown as cartoon, the small and large domains are colored in wheat and gray, respectively, and the cofactors FAD and NADP^+^ are shown as yellow and green sticks, respectively. Residues forming salt bridges that are unique to mFMO_20 are shown as sticks, hydrogen bonds between interacting residues are shown as teal-colored dashes, and oxygen and nitrogen atoms are colored red and blue, respectively. Native and mutated residues are colored blue and orange, respectively. The residues of salt bridges involving mutant residues are labeled. The three α-helices where the mutated residues are located are labeled in red. **B)** Chain A of the mFMO structure (PDB ID 2XVH) is visualized essentially as in A. Residues that form salt bridge pairs in both the native mFMO and mutant mFMO_20 are colored slate blue, and the lone pair which is unique to mFMO is colored cyan. **C)** The biological dimer of native mFMO is shown in B-factor putty representation. Red colors and large diameters of the tube indicate flexible regions with higher B-factors, in contrast to blue colors with small diameters, indicating well-ordered regions with lower B-factors. The location of the loop between β-strands 11 and 12 is indicated. Chain B is colored white. **D)** The biological dimer of mutant mFMO_20 is visualized as in C. The view of all structures has been rotated -60° around the y-axis compared to Figure 3.

**Table 4.** New polar interactions directly involving or induced by side chain atoms of mutated residues in mFMO_20. Hydrogen donor and acceptor atoms of the amino acid side chains are indicated in parenthesis, the distances between them are shown, and the structural location is indicated by secondary structure element.

| Hydrogen donor | Hydrogen acceptor | Distance (Å) | Structural location |  |
| --- | --- | --- | --- | --- |
|  |  |  | Donor | Acceptor |
| K300 (N<sup>ζ</sup>) | D351 (O<sup>δ2</sup>) | 3.0 | loop β18-β19 | α6 |
| K300 (N<sup>ζ</sup>) | N378D (O<sup>δ2</sup>) | 2.8 | loop β18-β19 | α7 |
| R356 (N<sup>ε</sup>/N<sup>η2</sup>) | M353Q (O<sup>ε1</sup>) | 2.8/2.7 | α6 | α6 |
| R356 (N<sup>η1</sup>) | L360E (O<sup>ε2</sup>) | 3.4 | α6 | α6 |
| K358R (N<sup>η1</sup>/N<sup>η2</sup>) | D374 (O<sup>δ2</sup>) | 3.0/2.8 | α6 | α7 |
| N394 (N<sup>δ2</sup>) | T370D (O<sup>δ1</sup>) | 3.4 | α8 | α7 |
| L398K (N<sup>ζ</sup>) | E366 (O<sup>ε2</sup>) | 3.1 | α8 | α7 |
| K401 (N<sup>ζ</sup>) | A365D (O<sup>δ2</sup>) | 2.7 | α8 | α7 |

The B-factor profile of mFMO_20 is different from that of native mFMO (Figure 4C and 4D), indicating differences in structural stability and flexibility. The region with the highest B-factor in mFMO_20 is the loop between β-strands 11 and 12, which is located at the entrance of the active site. In contrast, several loops in native mFMO display higher B-factors than that of the β11-β12 loop.

## Discussion

We have previously shown that mFMO can convert the malodorous TMA molecule into the odorless TMAO in a salmon protein hydrolysate (Goris et al., 2020). To make mFMO more suitable for such industrial applications, which typically takes place at temperatures ranging from 45 °C to 60 °C (Aspevik et al., 2016; Kristinsson and Rasco, 2000), we employed the PROSS algorithm to improve its thermal stability. All 7 mutant variants of mFMO were functional enzymes with increased thermostability (Table 3). The best variant was mFMO_20, which displayed the highest increase in temperature stability (Table 3). The crystal structure of mFMO_20 demonstrated that the overall structure of this mutant variant was highly similar to that of native mFMO, but also revealed new structural features that could explain the increased structural stability. The most striking new features were five salt bridges involving mutated residues, of which four formed stabilizing interhelical connecting bridges (Figure 4A). Although PROSS failed to correctly model the intrahelical salt bridge between L360E and R356 and predicted an intrahelical salt bridge between N290D and R292 that was not observed in the crystal structure, all four interhelical salt bridges were correctly modeled (Table 2), thus emphasizing the quality of the PROSS predictions. The only interhelical salt bridge that was present in all 7 PROSS models was the N378D-K300-D351 bridge, which connects α6 to α7 via the β18/β19-loop (Table 2). The N378D mutation leads to the replacement of a hydrogen bond between N378 and K300 in the native mFMO structure, by the stronger salt bridge between N378D and K300, and also induces K300 to form a salt bridge with D351, which is not present in the native structure. The fact that the largest increase in temperature stability from one variant to the next was observed going from native mFMO to mFMO_8, and that mFMO_8 only contains one interhelical salt bridge, may suggest that N378D is a key stabilizing mutation. The importance of the additional interhelical salt bridges is also corroborated by the fact that the observed gradual increase in temperature stability from mFMO_8 to mFMO_20 coincides with a gradual increase in the number of such bridges from 1 in mFMO_8 to 4 in mFMO_20 (Table 2). These results are in line with the recently published community-wide PROSS evaluation, where a correlation between the number of mutations and gain of thermal stability was observed (Peleg et al., 2021). However, despite introducing 4 and 8 more mutations, leading to two more salt bridges predicted to connect secondary structure elements in mFMO_24 (both between α7 and α8) and mFMO_28 (one between α7 and α8 and one between α6 and the β17/β18-loop), the temperature stability of these variants decreased slightly compared to mFMO_20. One possible explanation is that the intricate network of salt bridges in mFMO_24 and mFMO_28, which also involves direct bridges between mutated residues, are incorrectly predicted by PROSS.

In a previous effort to identify mutations that confer increased thermostability to mFMO, Lončar and colleagues used the FRESCO protocol to predict stabilizing single mutations (Lončar et al., 2019). The FRESCO analysis yielded 140 single mutant candidates that were expressed, purified, and screened for increased thermostability, and 14 of these displayed an apparent increase in melting temperature of >1 °C. The two mutations M15L and S23A were combined and the resulting mFMO variant had a 3 °C increase in melting temperature. Adding additional stabilizing single mutations did not further increase thermostability. In line with what we observed with mFMO_20, no major effects were observed on the kinetic parameters of the mFMO M15L/S23A variant. FRESCO has also been used to stabilize the *Rhodococcus* sp. HI-31 cyclohexanone monooxygenase (Fürst et al., 2019), which also belongs to the FMO family. In this case, half of the 128 screened single mutant variants had modest stabilizing effects. These were combined, using a shuffled library design strategy, into a variant carrying 8 mutations (M8B), which increased the unfolding temperature by 13°C. In contrast to the PROSS mFMO mutant variants, the FRESCO-derived mutations in neither M8B nor mFMO M15L/S23A appear to form new salt bridges.

We engineered mFMO to make it more suitable for industrial applications, such as removing TMA in salmon protein hydrolysates. The mFMO_20 variant was selected as the best candidate due to it being the most thermostable variant of the seven designs. In addition, the optimal pH for mFMO_20, pH 8.0, was slightly lower than that of native mFMO, pH 8.5, which may also confer an advantage in industrial applications (e.g., the pH of the salmon protein hydrolysate was 6.1). In fact, the optimal pH for all 7 mutant variants was between 7.5 and 8.0 (Table 3), but with no discernable correlation with changes in charge or pI. The optimal temperature of mFMO_20 did not increase compared to native mFMO. Still, this minor disadvantage of mFMO_20 was clearly outweighed by the beneficial properties of increased stability, when tested for its ability to convert TMA to TMAO in the salmon protein hydrolysate (Figure 2). At both 50 °C and 65 °C mFMO_20 was superior to native mFMO in removing TMA, eliminating 95% of TMA, and it also appeared to perform best at 30 °C. These results demonstrate that mFMO_20 is indeed more suitable for industrial applications than the native mFMO. However, there are still important hurdles that must be overcome before this Tmm enzyme can be incorporated into an industrial process, especially its dependence on the unstable and expensive cofactor NADPH. The fact that we and others have demonstrated that the Tmms can be engineered opens the possibility for cofactor engineering, which can be used to make the enzyme accept more cost-efficient cofactors. An alternative or complementary strategy is to bring down cost by regenerating the cofactor, e.g., by using glucose dehydrogenase (Mourelle-Insua et al., 2019).

The current work has demonstrated that the PROSS method successfully predicted mFMO variants with increased thermostability. All 7 variants proposed by PROSS, showed increased thermostability, with comparable properties to engineered FMO enzymes obtained after screening more than 100 single mutant variants followed by library shuffling or rational engineering (Fürst et al., 2019; Lončar et al., 2019). The mFMO_20 variant with its improved stability may be applicable for industrial use as it is because it can reduce the majority of TMA present in fish hydrolysates. It can also serve as an excellent starting point for rational engineering to further improve its catalytic efficiency or for cofactor engineering to make it accept more cost-efficient cofactors.

## Materials and Methods

### Protein stabilization mutagenesis using the PROSS webserver

The Protein Repair One-Stop Shop (PROSS) server (https://pross.weizmann.ac.il/) was used to predict variants of mFMO with increased stability (Goldenzweig et al., 2016). The mFMO structure (PDB ID: 2XVH) (Cho et al., 2011) was used as input and chain A was chosen as design target. To avoid mutating FAD- and NADPH-interacting residues, the small molecule ligands constraint was set to “FAD, NAP”, and to avoid mutating dimer interface residues, the interacting chains constraint was set to “B”. The multiple sequence alignment used as basis for the analyses was automatically generated by PROSS using the following default parameters: a minimal sequence identity of 30%, a maximum of 3,000 targets; and an E-value threshold of 0.0001. The PROSS server was accessed 7 October 2019. Salt bridges in the resulting structural models of mFMO variants were identified using the VMD (version 1.9.4) Salt Bridges Plugin (version 1.1) with default settings: the oxygen-nitrogen distance cut-off was 3.2 Å and the side-chain centers of mass cut-off was set to none. PyMOL (version 2.4, Schrödinger Inc., New York, NY, USA was used to confirm the predicted salt bridges involving mutated residues and to visualize the structural models. The multiple sequence alignment of native mFMO and the sequences of the PROSS mutants was visualized using a Python script and combined with the secondary structures of mFMO (PDB ID: 2XVH) visualized using ESPript (version 3.0).

### Molecular cloning of mFMO variants

The mFMO mutant variants predicted by PROSS were ordered as genes, codon-optimized for expression in *E. coli* and flanked by SapI sites, from TWIST Bioscience (San Francisco, CA, USA). Each mutation was introduced by changing the relevant codon of the native residue to one of the frequently used codons for the mutant residue in the *E. coli*, guided by the Codon Usage Database, and using the gene sequence encoding mFMO optimized for expression in *E. coli* as a starting point (Goris et al., 2020). The genes were subcloned into the C-terminal His-tag containing expression vector pBXC3H (p12) by fragment exchange cloning as previously described (Bjerga et al., 2016; Geertsma and Dutzler, 2011; Goris et al., 2020). Briefly, subcloning was performed using the *E. coli* MC1061 strain and Luria-Bertani (LB)-agar supplemented with ampicillin (100 μg/mL, Sigma-Aldrich, St. Louis, MO, USA) for selection, and the resulting plasmids, purified using the NucleoSpin plasmid kit (Macherey-Nagel, Düren, Germany), were confirmed by sequencing.

### Protein expression and purification

Native mFMO and mutant variants were expressed and purified essentially as previously described (Goris et al., 2020). Briefly, expression was performed using *E. coli* MC1061 cells in 100 mL LB-medium supplemented 100 μg/mL ampicillin at 20 °C for 16 hours after induction with 1% (w/v) L-arabinose. All purification steps were conducted at 4 °C. Cells were harvested by centrifugation, resuspended in lysis buffer, lysed by freeze thaw cycles and sonication, and cleared by centrifugation. The His-tagged mFMO variants were then purified from the cleared lysate using Ni-NTA resin. The buffer of the eluted protein was changed to 50 mM TrisHCl, pH 7.5, 100 mM NaCl using PD10 columns (GE Healthcare, Chicago, IL, USA). Finally, the protein was concentrated using protein concentrator columns (Thermo Fisher, Waltham, MA, USA) and stored with 10% glycerol at -20 °C until further use. Protein concentrations were measured using the Pierce^TM^ 660 nm Protein Assay Reagent (Thermo Fisher) with BSA as standard, and purity was assessed by polyacrylamide gel electrophoresis (SDS-PAGE).

### Peptide Mass Fingerprinting by Matrix-Assisted Laser Desorption/Ionization-Time-of-Flight/Time-Of-Flight (MALDI-TOF/TOF)

MALDI-TOF/TOF analysis in-solution of purified protein samples was performed as previously described (Santiago et al., 2018). The confidence interval for protein identification was set to ≥95% (*p* < 0.05) and only peptides with an individual ion score above the identity threshold were considered correctly identified. The analysis was performed at the Unidad de Proteómica, Centro Nacional de Biotecnología (CNB-CSIC), Madrid, Spain (analysis ID 3408).

### Enzyme activity assay

Enzyme activity towards TMA was assessed by monitoring the consumption of the cofactor NADPH as previously described (Goris et al., 2020). Briefly, the assay was performed in 96-well microtiter plates, using reaction buffer (50 mM Tris-HCl pH 8.0, 100 mM NaCl) supplemented with 0.5 mM NADPH (Merck, Rahway, NJ, USA/Sigma-Aldrich) and 0.01-0.02 mg/mL enzyme. The reactions were initiated by adding 1 mM TMA (Sigma-Aldrich), and consumption of NADPH was measured by continuously monitoring absorbance at 340 nm over 30 minutes using an Epoch^TM^ Microplate Spectrophotometer (Biotek, Winooski, VT, USA). Initial reaction rates were determined from linear fits of the absorbance versus time corrected for blank. Assays were performed in triplicates at 22 °C, unless otherwise stated. One unit (U) of enzyme activity was defined as the number of enzymes required to transform 1 μmol substrate in 1 minute under the stated assay conditions and using the extinction coefficient ε_340_ = 6.22 mM^-1^ cm^-1^ for NADPH.

### pH optimum

pH optimum was measured using the enzyme activity assay as described above, in a three-component buffer (100 mM sodium acetate, 50 mM Bis-Tris and 50 mM Tris) that was pH adjusted using 100% acetic acid. pH dependence was investigated between pH 6.0 and 9.0, with increments of 0.5, and the enzymes were incubated for 2 h in the appropriate buffer at a fixed temperature of 22 °C before measuring residual enzymatic activity by adding NADPH and TMA. The experiment was performed on one biological replicate (enzyme preparation) with technical triplicates.

### Temperature optimum

Optimal temperature was measured using the enzyme activity assay described above in 1 mL cuvettes using a Cary 60 UV-Vis spectrophotometer (Agilent Technologies, Santa Clara, CA, USA) at 22 °C and temperatures from 30 °C to 50 °C with 5 °C increments using a circulating water bath. The enzymes were diluted to 0.01 mg/mL in 1 mL buffer (50 mM Tris-HCl, pH 8.0, 100 mM NaCl) preheated in a water bath to the corresponding temperature and incubated for 1 minute together with 0.5 mM NADPH before initiating the reaction with 1 mM TMA. Two biological replicates (enzyme preparations) were tested, each with 1 to 3 technical replicates resulting in at least three replicates per temperature step for each enzyme. No reliable measurements were obtained for native mFMO at 50 °C.

### Temperature stability

To assess temperature stability, freshly purified enzymes were diluted to 0.5 mg/mL in reaction buffer (50 mM Tris-HCl pH 7.5, 100 mM NaCl) and initial enzyme activity was measured as described above. The enzymes were subsequently incubated at temperatures from 30 °C to 54 °C for one hour using a PCR thermocycler machine (Bio-Rad Laboratories, Hercules, CA, USA), followed by cooling for 10 minutes at 4 °C, incubation at RT for 10 minutes, and centrifugation for 2 min using a tabletop centrifuge. The residual enzyme activity was measured as described above and recorded as relative to the initial activity. The temperature where half of the initial activity was lost was determined by four-parameter logistic regression using Prism 9 (GraphPad Software, San Diego, CA, USA). The experiment was performed three times using two different enzyme preparations, each time with three technical triplicates.

### Enzyme steady state kinetics

To determine the *K*_M_ for TMA mFMO and mFMO_20 were purified essentially as described above and flash frozen in 20% glycerol. 50 or 25 pmol of enzyme was diluted in 880 µl reaction buffer (50 mM tris-HCl, pH 8.0) in a cuvette, and mixed with 20 µL 10 mM NADPH (final concentration: 200 µM). The enzyme and cofactor were incubated for 2 minutes at room temperature before 50 µl TMA (final concentration between 1-500 µM) was added to the cuvette. This was mixed thoroughly, and the absorbance decrease at 340 nm was immediately measured for 2 minutes in a Cary 60 UV-Vis spectrophotometer (Agilent Technologies) at 23 °C to obtain the initial reaction rate of the enzymes (V_0_), defined as the change in absorbance per minute in the linear part of the curve. To calculate the product formation per enzyme (µmol product min^-1^ µmol enzyme^-1^), the NADPH conversion rate was calculated from the absorbance change assuming a molar extinction coefficient of NADPH of 6220 M^-1^ cm^-1^. The product formation per enzyme was plotted against the TMA concentration, and *K*_M_ and k_cat_ were calculated using nonlinear regression in Prism 9. Each enzyme was expressed and purified in two biological replicates, and for each biological replicate two replicate measurements were made and averaged.

### CD spectroscopy

Melting temperatures of the mFMO variants were recorded by circular dichroism (CD) spectrometry essentially as previously described (Goris et al., 2020). Briefly, the enzymes were diluted to 0.7-0.8 mg/mL in reaction buffer (50 mM Tris-HCl pH 7.5, 100 mM NaCl) and denaturation was monitored, using 0.1-cm path length quartz cuvettes, at 220 nm between 10 °C and 95 °C at a rate of 30 °C per hour using a Jasco J-720 spectropolarimeter (Japan Spectroscopic Corporation, JASCO, Tokyo, Japan) equipped with a Peltier temperature controller. The melting temperature was calculated by fitting the ellipticity (mdeg) at 220 nm for each of the different temperatures using four-parameter logistic regression using Prism 9 (GraphPad Software).

### Enzymatic conversion of TMA to TMAO in salmon protein hydrolysates

Enzymatic conversion of TMA present in the salmon protein hydrolysate to TMAO was performed essentially as described previously (Goris et al., 2020). The salmon protein hydrolysate (64.4% dry weight) was produced from fresh salmon by-products by protease treatment and provided by the Biomega Group (Skogsvåg, Norway). The viscous salmon protein hydrolysate was diluted 1:5 (wt/vol) in ultrapure water, followed by sonication in an ultrasonic water bath (J.P. Selecta, S.A., Barcelona, Spain) at 50 Hz for 5 min, vortexing for 5 min, and centrifugation at 16,000 × *g* for 10 min. The resulting supernatant (pH 6.10) was supplemented with 0.50 mM NADPH and 10 ng/mL mFMO or mFMO_20 enzyme and incubated for 1 h at 30 °C, 50 °C and 65 °C. The enzymatic reaction was stopped by diluting 10 times with methanol. All reactions were performed with two technical triplicates using one enzyme preparation and included control samples without enzyme. The TMA levels in the samples were determined using Ultra High Performance Liquid Chromatography (UHPLC) with EVOQ Elite Triple Quadrupole Mass Spectrometer (Bruker, Billerica, MA, USA), each replicate measured two times. Prior to the analysis the samples were diluted 1:1000 by mixing with methanol and subsequently vortexed for 1 min. TMA and TMAO standards were prepared in water to a final concentration of 25, 50, 100, 200 and 250 ppb. A liquid chromatography system consisting of a degasser, a binary pump, and an auto-sampler (at 4 °C) was used. Samples were applied to a column (Ace Excel 3, C18-Amida, 3µm, 150 x 4.6mm ID; Advanced Chromatography Technologies Ltd., Reading, UK), which was maintained at 40 °C during the analysis. The system was operated at a flow rate of 0.5 mL/min with solvent A (H_2_O containing 0.1% formic acid) and solvent B (methanol). The system was held at 2% B for 7 min of total analysis time. Data were collected in positive electrospray ionization (ESI) mode using Q-TOF (model Agilent 6120; Agilent Technologies). The spray voltage was 5000 V, the cone temperature 350 °C, the cone gas flow 40 L/h, the heated probe temperature 400 °C, the probe gas flow 50 L/h, and the nebulizer gas flow 60 L/h. The experiment was performed twice with technical duplicates. The analyses were performed at the Servicio Interdepartamental de Investigación (SIDI) from the Autonomous University of Madrid (analyses ID 200-01807, 200-01761, and 200-01629).

### Crystallization of mFMO_20

The complex of mFMO_20 with NADP^+^ and FAD was obtained by co-crystallization assays incubating 5.13 mg/ml protein in 20 mM Tris pH 8, 150 mM NaCl, 1 mM DTT with 1 mM NADPH during 25 min at 4 °C. Initial crystallization conditions were explored by a NanoDrop robot (Innovadyne Technologies, Santa Rosa, CA, USA) and the commercial screen Index (Hampton Research, Aliso Viejo, CA, USA). Yellow prism bar shaped crystals were grown after two months by adding 250 nl of the protein mixture to 250 nl of precipitant solution (2 M (NH_4_)_2_SO_4_, 0.1 M Bis-Tris pH 6.5). For data collection, crystals were transferred to a cryoprotectant solution consisting of 2.2 M (NH_4_)_2_SO_4_, 0.1 M Bis-Tris pH 6.5 and 23% (v/v) glycerol, before being cooled in liquid nitrogen.

### Data collection and structure determination

Diffraction data were collected using synchrotron radiation on the XALOC beamline at ALBA (Cerdanyola del Vallés, Spain). Diffraction images were processed with XDS (Kabsch, 2010) and merged using AIMLESS from the CCP4 package (Evans and Murshudov, 2013). The crystals were indexed in the C222_1_ space group, with two molecules in the asymmetric unit and 44% solvent content within the cell. The structure of mFMO_20 complexed with NADPH was solved by Molecular Replacement with MOLREP (Vagin and Teplyakov, 2010) using the coordinates from the wild type as template (PDB ID: 2XVE). Crystallographic refinement was performed using the program REFMAC (Murshudov et al., 1997) within the CCP4 suite with local non-crystallographic symmetry (NCS). A summary of the data collection and refinement statistics is found in Table S1. Free R-factor was calculated using a subset of 5% randomly selected structure-factor amplitudes that were excluded from automated refinement. At the later stages, ligands were manually built into the electron density maps with COOT (Emsley et al., 2010) and water molecules were included in the model, and combined with more rounds of restrained refinement. The figures were generated with PyMOL. The structure is available with PDB ID 8B2D.

### Identification of polar interactions

To perform an inclusive identification of salt bridges in both the native mFMO (PDB ID: 2XVH) and mFMO_20 structures, MolProbity (Williams et al., 2018) was used to add hydrogen atoms and optimize polar contacts by side-chain flip correction of Asn, Gln, and His residues. Residues that were flipped in both structures were Q17, Q40, N48, H164, and N282, additional residues flipped in mFMO were Q25, Q143, and N290, and residues flipped only in mFMO_20 were H128, M353Q, Q377, and H414. Chain A from these optimized structures were used to identify salt bridges with the VMD Salt Bridges Plugin (as described above). PyMOL was used to confirm the predicted salt bridges, to identify additional hydrogen bonds involving mutated residues, and to visualize the structural models.

## Acknowledgments

Authors would like to acknowledge Sergio Ciordia at the Proteomic Facilities from the CSIC-CNB (that belongs to ProteoRed, PRB2-ISCIII, supported by Grant PT13/0001) for the MALDI-TOF/TOF analyses, Rosa Sedano and Eva Martín at the Servicio Interdepartamental de Investigación (SIDI) from the Autonomous University of Madrid for the UHPLC-MS analyses, and Ruth Matesanz from the CSIC-CIB for her support for CD analyses. G.E.K.B gratefully acknowledge funding from the Research Council of Norway with an additional mobility grant to M.G. (RCN grant number 280737) as well as funding from the European Union’s Horizon 2020 research and innovation programme under grant agreement No 101000607. M.F. and J-S.A. acknowledge the European Union’s Horizon 2020 research and innovation programme under grant agreement No 101000327, and the financial support under Grants PID2020-112758RB-I00 (M.F.), PDC2021-121534-I00 (M.F.) and PID2019-105838RB-C33 (J.S-A.) from the Ministerio de Ciencia e Innovación, Agencia Estatal de Investigación (AEI) (Digital Object Identifier 10.13039/501100011033), Fondo Europeo de Desarrollo Regional (FEDER) and the European Union (“NextGenerationEU/PRTR”), and Grant 2020AEP061 (M.F.) from the Agencia Estatal CSIC. J-S.A. and I.C-R., thank the staff of the Synchrotron Radiation Source at Alba (Barcelona, Spain) for assistance at BL13-XALOC beamline. G.E.K.B. and M.G. would like to thank M.F. and CSIC for hosting the mobility stay. Authors thank Bjørn Liaset at the Biomega Group for providing salmon protein hydrolysate.

## Author contributions

G.E.K.B. and P.P. performed the PROSS analysis. M.G. planned experiments, cloned and purified all constructs, conducted biochemical experiments and analyzed all data. I.C.R. and J.S-A. conducted crystallography and structural analysis. R.R. conducted kinetic analyses. D.A. conducted CD experiments. M.F. supervised and prepared all mass spectrometry experiments together with M.G. G.E.K.B. secured the main funding, conceptualized the study, and supervised the project together with P.P. and M.F. M.G. and P.P. wrote the paper with input from I.C.R., J.S-A., R.R., M.F., and G.E.K.B. All authors read and revised the manuscript.

## Competing interests

The authors declare no conflict of interest.

## Data Availability

Coordinates and structure factors have been deposited in the Protein Data Bank under PDB accession code 8B2D.

## Notes

### Competing Interest Statement

The authors have declared no competing interest.

### Summary of Updates

Supplemental data is added

