## Supplemental data for "Increased thermostability of an engineered flavin-containing monooxygenase to remediate trimethylamine in fish protein hydrolysates"

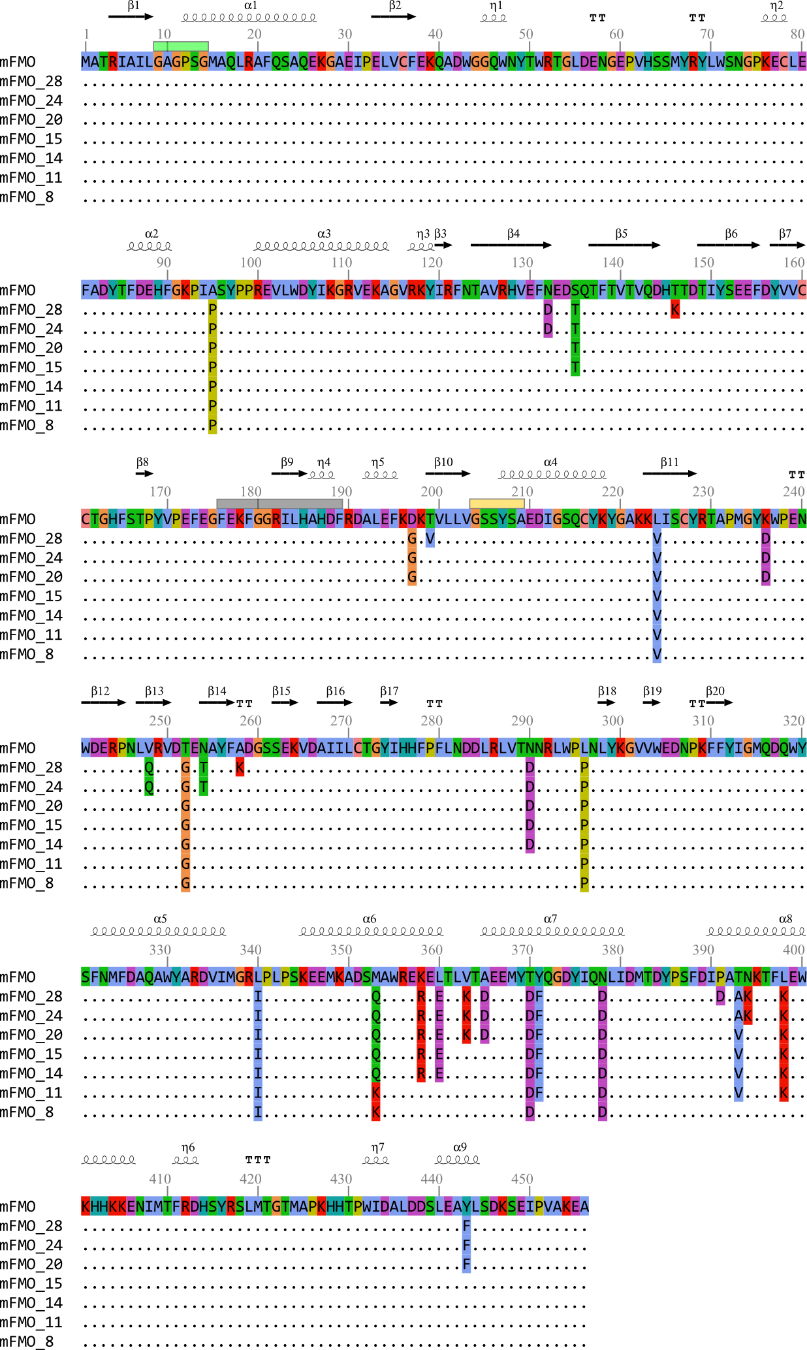

**Figure S1. Multiple sequence alignment of mFMO and mutant variants.** The sequences of the mutant variants generated using PROSS are listed in descending order according to number of mutations, which are indicated by the number after the underscore in the sequence names. Amino acid residues of the native mFMO sequence and mutant residues are shown in Clustal colors, and variant sequence residues identical to those of native mFMO are indicated with dots. Secondary structure elements of native mFMO are indicated above the alignment (PDB:2XVH). β-strands 8 and 17 were missing in the PDB file and were added manually (Cho et al 2011). The boxes above the alignment indicates the following sequence motifs: NADPH binding motif (green), FMO signature motif (grey), and FAD binding motif (yellow).

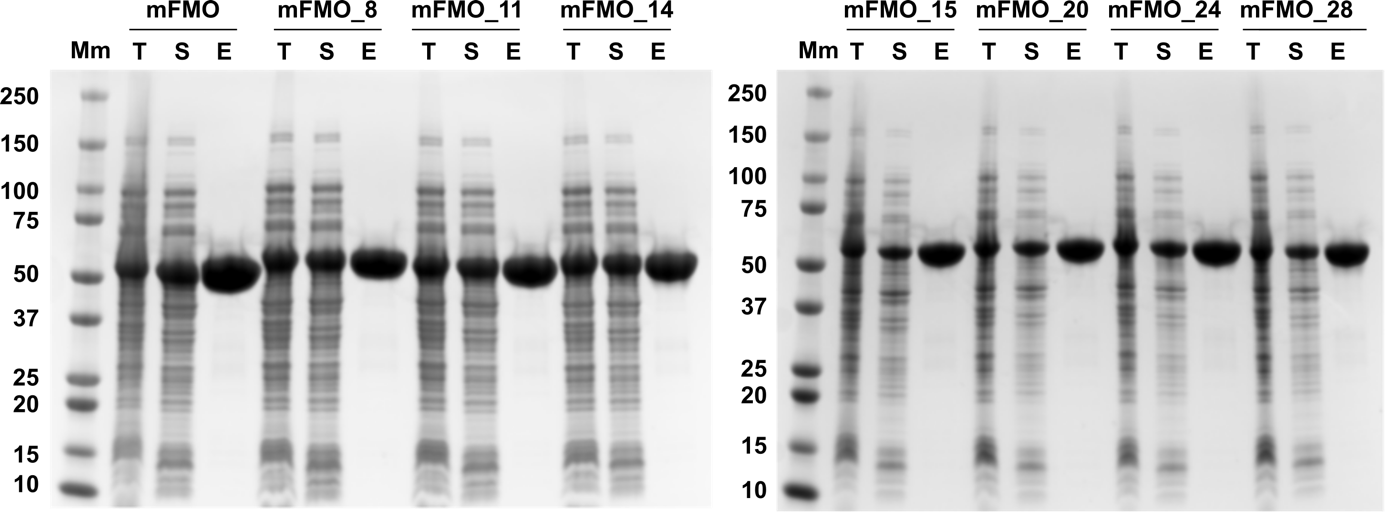

**Figure S2. Expression and purification mFMO and mFMO mutant variants.** The enzyme variants were expressed in E. coli and purified by Ni-NTA affinity chromatography. The figure shows SDS-PAGE gels documenting the total protein (T) after expression, the soluble proteins (S) present in the clarified lysate, and the eluate after Ni-NTA purification (E). Molecular masses (kDa) of the protein standard molecular marker (Mm) are shown to the left of each gel.

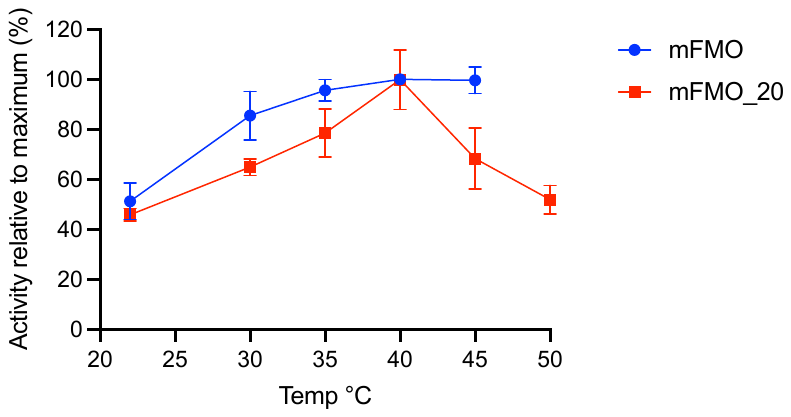

**Figure S3. Temperature optimum.** Specific activity (U/mg) of mFMO (blue) and mFMO_20 (red) against TMA at pH 8.0 was measured across temperature (22 to 50 °C) by NADPH consumption (each with three technical replicates). Activity is shown relative to maximal activity (100%) for each enzyme. One unit (U) of enzyme activity was defined as the amount of enzyme required to transform 1 μmol TMA in 1 min under the assay conditions using the reported NADPH extinction coefficient (ε_340_ = 622 mM^-1^ cm^-1^). For mFMO, no datapoints could be collected at 50 °C.

**Table S1. Crystallographic statistics of mFMO_20**

| **Crystal data** |  |
| --- | --- |
| Space group | C222_1_ |
| Unit cell parameters |  |
| a (Å) | 125.61 |
| b (Å) | 130.36 |
| c (Å) | 115.65 |
| **Data collection** |  |
| Beamline | XALOC(ALBA) |
| Temperature (K) | 100 |
| Wavelength (Å) | 0.979264 |
| Resolution (Å) | 45.23-1.62 (1.65-1.62) |
| **Data processing** |  |
| Total reflections | 1304601 (65330) |
| Unique reflections | 120046 (5926) |
| Multiplicity | 10.9 (11.0) |
| Completeness (%) | 100.0 (100.0) |
| Mean *I*/σ (*I*) | 20.7 (4.1) |
| R_merge_^†^ (%) | 6.5 (70.1) |
| R_pim_^††^ (%) | 2.1 (22.0) |
| Molecules per ASU | 2 |
| **Refinement** |  |
| R_work_ / R_free_*^†††^* (%) | 16.2/18.0 |
| Nº of atoms/average B (Å^2^) | 8111/23.16 |
| Macromolecule | 7352/22.76 |
| Ligands | 264/24.91 |
| Solvent | 495/28.25 |
| Ramachandran plot (%) |  |
| Favoured | 95.0 |
| Outliers | 0.2 |
| **RMS deviations** |  |
| Bonds (Å) | 0.008 |
| Angles (°) | 1.434 |
| **PDB accession code** | 8B2D |

^†^Rmerge = ∑hkl ∑i | Ii(hkl) – [I(hkl)]| / ∑hkl ∑i Ii(hkl), where Ii(hkl) is the ith measurement of reflection hkl and [I(hkl)] is the weighted mean of all measurements.

^††^Rpim = ∑hkl [1/(N - 1)] 1/2 ∑i | Ii(hkl) – [I(hkl)]| / ∑hkl ∑i Ii(hkl), where N is the redundancy for the hkl reflection.

^†††^Rwork / Rfree = ∑hkl | Fo – Fc | / ∑hkl | Fo |, where Fc is the calculated and Fo is the observed structure factor amplitude of reflection hkl for the working / free (5%) set, respectively.
